# Is photosynthetic enhancement sustained through three years of elevated CO_2_ exposure in 175-year old *Quercus robur*?

**DOI:** 10.1101/2020.12.16.416255

**Authors:** A Gardner, DS Ellsworth, KY Crous, J Pritchard, AR MacKenzie

## Abstract

Current carbon cycle models attribute rising atmospheric CO_2_ as the major driver of the increased terrestrial carbon sink, but with substantial uncertainties. The photosynthetic response of trees to elevated atmospheric CO_2_ is a necessary step, but not the only one, for sustaining the terrestrial carbon uptake, but can vary diurnally, seasonally and with duration of CO_2_ exposure. Hence we sought to quantify the photosynthetic response of the canopy-dominant species, *Quercus robur*, in a mature deciduous forest to elevated CO_2_ (eCO_2_) (+150 μmol mol^-1^ CO_2_) over the first three years of a long-term free air CO_2_ enrichment facility at the Birmingham Institute of Forest Research in central England (BIFoR FACE). Over three thousand measurements of leaf gas exchange and related biochemical parameters were conducted in the upper canopy to assess the diurnal and seasonal responses of photosynthesis during the 2^nd^ and 3^rd^ year of eCO_2_ exposure. Measurements of photosynthetic capacity via biochemical parameters, derived from CO_2_ response curves, (*V*_cmax_ and *J*_max_) together with leaf nitrogen concentrations from the pre-treatment year to the 3 ^rd^ year of eCO_2_ exposure, were examined. We hypothesized an initial enhancement in light-saturated net photosynthetic rates (A_sat_) with CO_2_ enrichment of ≈37% based on theory but also expected photosynthetic capacity would fall over the duration of the study. Over the three-year period, A_sat_ of upper-canopy leaves was 33 ± 8 % higher (mean and standard error) in trees grown in eCO_2_ compared with ambient CO_2_ (aCO_2_), and photosynthetic enhancement decreased with decreasing light. There were no significant effects of CO_2_ treatment on *V_cmax_* or *J*_max_, nor leaf nitrogen. Our results suggest that mature *Q. robur* may exhibit a sustained, positive response to eCO_2_ without photosynthetic downregulation, suggesting that, with adequate nutrients, there will be sustained enhancement in C assimilated by these mature trees. Further research will be required to understand the location and role of the additionally assimilated carbon.

## Introduction

Forest ecosystems cover about 30% of the Earth’s land surface, representing ~50% of terrestrially stored carbon and account for close to 60% of total terrestrial CO_2_ fluxes in the global carbon cycle (Luyssaert et al., 2008; Pan et al., 2011). The continual rise in atmospheric CO_2_, overwhelmingly due to anthropogenic activity (Friedlingstein et al., 2019), increases the need to understand the terrestrial carbon feedbacks of forests in the global carbon cycle. As the foundational driver of the carbon cycle of forests (e.g. Bonan, 2008), the photosynthetic response to changing atmospheric CO_2_ is a necessary process for forests to act as long-standing carbon stores with relatively long-lived carbon (C) pools such as wood (Körner, 2017) and soil (Ostle et al., 2009). The amount of forest C-uptake in the future, and subsequent C sequestration, will be crucial determinants of future atmospheric CO_2_ concentrations. So, quantifying the photosynthetic response under elevated CO_2_ (eCO_2_), especially for mature trees, is critical to understanding the carbon uptake of forests under changing atmospheric composition.

It has been widely observed that eCO_2_ can have a stimulatory effect on plant photosynthesis, known as photosynthetic enhancement, at least in the short-term (weeks to months) with adequate nutrient and water availability permitting (Brodribb et al., 2020). Long-term (years to decades) photosynthetic responses to eCO_2_ are less well understood and lower-than-expected responses have been observed (Ainsworth & Long, 2005; Ellsworth et al., 2017). Note that, even in studies that report sustained and/or strong stimulation of photosynthesis under eCO_2_, the additionally assimilated C does not necessarily translate into increased growth stimulation (Bader et al., 2013; Sigurdsson et al., 2013).

The photosynthetic process and photosynthetic response to eCO_2_ is sensitive to changes in environmental variables such as temperature, light, water and availability of nutrients. For example, net photosynthesis (A_net_) is expected to increase with exposure to eCO_2_, with greatest photosynthetic enhancement expected at maximum photon flux density (*Q*) if Rubisco carboxylation is limiting (Sage et al., 2008). Decreases in A_net_ have been commonly associated with limitations in water and nutrient availability (Ainsworth & Rogers, 2007; Nowak et al., 2004). For example, water availability has been found to increase the magnitude of eCO_2_-induced photosynthetic enhancement in drier years (Ellsworth et al., 2012; Nowak et al., 2004). Thus, interannual differences in eCO_2_–induced photosynthetic enhancement are to be expected as environmental conditions vary. Understanding the photosynthetic response to eCO_2_ under different, real-world, environmental conditions provides information essential, but not in itself sufficient, for modelling forest productivity (Jiang et al., 2020), and predicting carbon-climate feedbacks (e.g., Cox et al., 2013; Jones et al., 2016).

Despite a significant body of research on the photosynthetic response to eCO_2_ in tree seedlings and saplings (as reviewed in Ainsworth & Long, 2005; Medlyn et al., 1999), fewer studies address the long-term (> 1 year) photosynthetic responses in mature plantation trees (Crous et al., 2008; Liberloo et al., 2007; Uddling et al., 2009; Warren et al., 2015) and very few in mature forest-grown trees (Bader et al., 2010; Ellsworth et al., 2017; Klein et al., 2016). Currently, the dynamic vegetation components of Earth System models, which diagnose vegetation responses to environmental change, have commonly been constructed using data from eCO_2_ experiments on young and/or plantation grown trees (Piao et al., 2013). Yet, it is difficult to compare, generalise, and scale results from young trees in their exponential growth phase to the response of closed-canopy mature forests (Norby et al., 2016). For example, previous work from a long-term natural experiment found CO_2_ stimulation declined with tree age in *Quercus ilex* (Hättenschwiler et al., 1997). Therefore, it is plausible that model projections are currently overestimating the photosynthetic responses of mature forests and, thence, the ‘CO_2_ fertilisation’ effect (Zhu et al., 2016). Consequently, uncertainty remains as to the magnitude of, and environmental constraints on, photosynthetic enhancement under eCO_2_ in large, long standing carbon stores such as mature forests (Jiang et al., 2020; Norby et al., 2016).

Free-air CO_2_ enrichment (FACE) facilities are valuable to understand system-level responses to eCO_2_ (Ainsworth & Long, 2005; Terrer et al., 2019) particularly in forests (Medlyn et al., 2015; Norby et al., 2016). The development of 2^nd^ generation forest FACE experiments focuses on tall, mature trees grown in their own forest soil (Hart et al., 2020). To date, forest FACE experiments have observed photosynthetic enhancements ranging from 30-60%, depending on tree species and environmental factors (as reviewed in Ainsworth & Rogers, 2007; Nowak et al., 2004). Of the few studies on closed-canopy dominant tree species, smaller photosynthetic enhancement to eCO_2_ have been observed (19 to 49%) than in studies conducted on younger trees (Crous et al., 2008; Liberloo et al., 2007; Sholtis et al., 2004), but the reasons behind this smaller response remain unclear.

There is evidence of a reduction in photosynthetic activity after long-term eCO_2_ exposure, known as photosynthetic downregulation (Ainsworth et al., 2004; Crous & Ellsworth, 2004), but downregulation is not always observed (Curtis & Wang, 1998; Herrick & Thomas, 2001). Commonly photosynthetic downregulation under eCO_2_ exposure is the result of decreases, either directly or indirectly, in Rubisco carboxylation (*V*_cmax_) (Feng et al., 2015; Wujeska-Klause et al., 2019b). However, the stimulatory effect of photosynthesis under eCO_2_ may be reduced but is usually not completely removed (Leakey et al., 2009; Wujeska-Klause et al., 2019). Photosynthetic downregulation has largely been observed in young plants (Leakey et al., 2009), with some downregulation observed in two aggrading plantation forests (Crous et al., 2008; Warren et al., 2015), commonly as a result of insufficient soil nitrogen supply (Luo et al., 2004). However, photosynthetic downregulation has largely not been observed in mature forests (Bader et al., 2010) and therefore open questions remain concerning the frequency and magnitude of photosynthetic downregulation under eCO_2_ exposure in mature forests.

To understand the photosynthetic responses in mature temperate deciduous forests, we evaluated the photosynthetic enhancement and potential downregulation in ca. 175-year old canopy dominant trees of *Quercus robur* L. exposed to elevated CO_2_ for three years. Considering that forest FACE experiments aim to operate for 10 years or more, we refer to these CO_2_ responses as ‘early’ (Griffin et al., 2000). This study is amongst the oldest trees that have ever been examined under elevated CO_2_. To assess the photosynthetic enhancement of the trees on daily and interannual timeframes, measurements of gas exchange and leaf biochemistry were measured in the upper oak canopy over four growing seasons, that included one pre-treatment year (2015) and three treatment years (2016 to 2019). Our aims were to quantify the photosynthetic response to eCO_2_ (i.e., ambient +150 μmol mol^-1^) for mature *Q. robur* and how light level influences this response, to determine whether photosynthetic downregulation under eCO_2_ occurred and to establish whether the relationship between leaf N and photosynthetic capacity changed in eCO_2_. We hypothesized that net photosynthetic gas exchange, A_net_, will significantly increase with eCO_2_ and light levels (*Q*). The greatest enhancement was expected with the highest light levels, as a result of reduced limitations in the light dependent reaction of photosynthesis, and that photosynthetic enhancement would be ≈37% following theory and reasoning in Nowak et al. (2004)(see also Supplemental Appendix 1. below). We also hypothesized that leaf nitrogen (N) will be reduced under elevated CO_2_ and that photosynthetic downregulation will be observed under eCO_2_ as a result of reduced leaf N and/or a decline in either the maximum rate of photosynthetic Rubisco carboxylation (*V*_cmax_, μmol m^-2^ s^-1^); and the maximum rate of photosynthetic electron transport (*J*_max_, μmol m^-2^ s^-1^), or both (Griffin et al., 2000).

## Methods and materials

### Site description

This study was conducted at the Birmingham Institute of Forest Research (BIFoR) Free Air CO_2_ Enrichment (FACE) facility located in Staffordshire (52.801°N, 2.301°W), United Kingdom. The BIFoR FACE facility is a ’2^nd^ generation’ Forest FACE facility, extending the scope of 1^st^ generation facilities; (see Norby et al., 2016), situated within 19 ha of mature northern temperate broadleaf deciduous woodland having a canopy height of 24-26 m. The woodland consists of an overstorey canopy dominated by English oak (*Quercus robur* L.) and a dense understorey comprising mostly of hazel coppice (*Corylus avellana* L.), sycamore (*Acer pseudoplatanus* L.), and hawthorn (*Crataegus monogyna* Jacq.). *Q. robur* (commonly known as pendunculate oak, European oak or English oak) is a common broadleaf species geographically widespread across Europe where it is both economically important and ecologically significant for many biota (Eaton et al., 2016; Mölder et al., 2019). The site was planted with the existing oak standards in the late 1840s and has been largely unmanaged for the past 30 to 40 years. Like most established forest of the temperate zone, the BIFoR FACE forest is under-managed.

The study site is situated within the temperature-rainfall climate space occupied by temperate forest (Jiang et al., 2020; Sommerfeld et al., 2018) and is characterized by cool wet winters and warm dry summers with a frost-free growing season from April to October. The mean January and July temperatures were 4 and 17 °C, respectively, and the average annual precipitation for the region is 720 mm (650 mm, 669 mm, 646 mm and 818 mm, in 2015, 2017, 2018 and 2019, respectively, when the study was conducted; see Figure 1.). The total N deposition load at the BIFoR FACE site is ~22 Kg N/ha/year (estimate provided by S. Tomlinson at the Centre for Ecology and Hydrology, Edinburgh, UK)(Mackenzie et al., 2021), representing around 15% of the total nitrogen nutrition of temperate deciduous trees (Rennenberg & Dannenmann, 2015).

**Figure 1.**
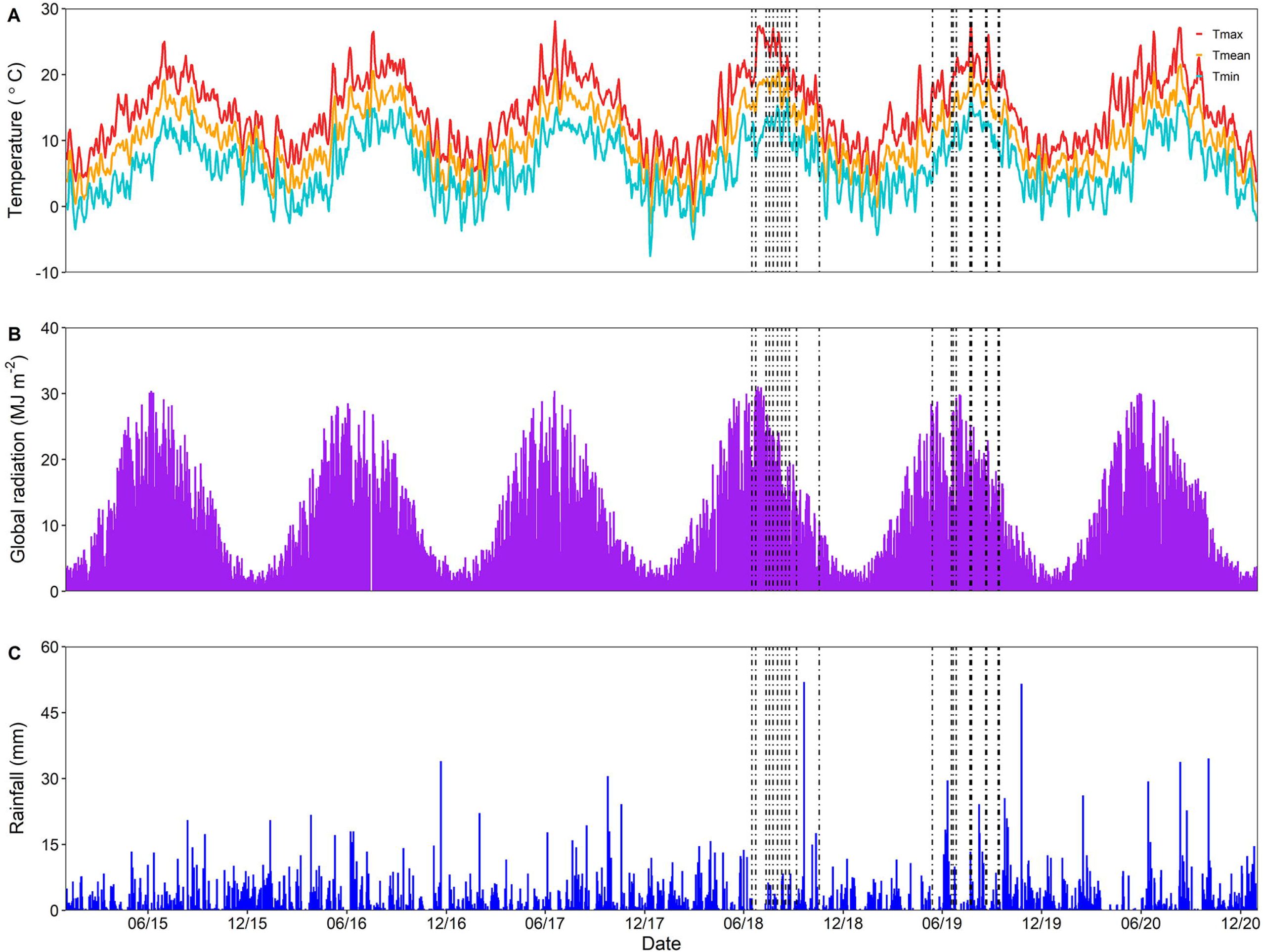
Time series showing the daily meteorological data at the BIFoR FACE facility covering the period of 01-01-2015 to 01-01-2021. Subplots are: A) maximum, (red), mean (orange) and minimum (blue) daily air temperatures (°C), B) global downwelling solar radiation (MJ m^-2^) and C) total daily precipitation (mm). Vertical dashed lines indicate diurnal sampling days. Clusters of sampling days occurred because different plots were sampled on different days in the same seasonal timeframe. Meteorological data is from RAF Shawbury, located 20 miles west of the BIFoR FACE facility, retrieved from the UK Met Office (https://www.metoffice.gov.uk/research/climate/maps-and-data/historic-station-data).

BIFoR FACE consists of nine approximately circular experimental plots of woodland 30 m in diameter (Hart et al., 2020). Only the six plots with infrastructure were considered in the present study. Each ‘infrastructure plot’ is encircled by steel towers constructed individually to reach 2 m above the local canopy-top height. The facility uses a paired-plot design (Hart et al., 2020): three replicate plots at either ambient CO_2_(aCO_2_) (ca. 405 μmol mol^-1^) and three plots supplied with CO_2_ enriched air, termed elevated CO_2_ plots (eCO_2_). The latter plots were operated such that they achieved a target of +150 μmol mol^-1^ above the minimum measured in the ambient plots (i.e. concentrations in the elevated plots ca. 555 μmol mol^-1^) as five-minute rolling averages (Hart et al., 2020)(Supplementary Figure 1). Elevated CO_2_ is added from dawn (solar zenith angle, sza = −6.5°) to dusk (sza = −6.5°) throughout the growing season. Daytime exposure to eCO_2_ was almost continuous throughout the growing season (Hart et al., 2020), with exceptions if the 15-minute average wind speed was greater than 8 m s^-1^, or when canopy-top, 1-min average, air temperature was < 4°C. In the latter case, gas release was resumed when the air temperature was ≥ 5°C. The CO_2_ fumigation thresholds for wind speed and temperature were selected because of the high cost of maintaining elevated CO_2_ and the insignificant uptake of carbon under these conditions, respectively. The operation of the FACE system and statistical performance in terms of meeting the target CO_2_ concentration in time and space have been described in Hart et al. (2020).

In each plot, canopy access was gained through a custom-built canopy access system (CAS) (Total Access Ltd., UK) that was installed from the central towers with canopy measurements made from a rigged rope access system (Supplementary Figure 2.). This facilitated *in situ* gas exchange measurements by allowing access to the upper oak canopy. The hoisting system comprises of an electric winch (Harken Power Seat Compact) that lifts a harnessed (Petzl AVAO BOD 5 point harness) user vertically through the air at a predetermined fixed point to a maximum canopy height of 25 m. The system required operation from the ground by trained staff and the user is seated in a Boatswain’s chair. One oak tree per plot was accessible using the CAS system as set up during this study, and all gas exchange measurements were made on unshaded leaves within the top two meters of each tree canopy on dominant trees.

For this study, the sample size used throughout the study (n=3) represents the number of replicate experimental plots at BIFoR FACE and includes within-tree replicates that were averaged per plot before analysis. All the three replicates were sampled for the majority of campaigns, except for September 2018 and June 2019 where replicates were reduced to two due to logistic constraints, weather, and safe tree access.

### Gas exchange measurements

All gas exchange measurements were conducted *in situ* on upper canopy oak leaves using either a Li-6400XT or Li-6800 portable photosynthesis system (LiCOR, Lincoln, NE, USA) to quantify photosynthetic performance at BIFoR FACE. Measurement campaigns focussed on two different types of measurements: i) instantaneous diurnal measurements, at prevailing environmental conditions (2018 and 2019), and ii) net assimilation rate-intercellular CO_2_ concentration (*A-–C*_i_) measurements (includes pre-treatment, 2015; 1^st^ year, 2017; and 3^rd^ year, 2019, of CO_2_ fumigation). Measurements were conducted in all six experimental plots with infrastructure, on one chosen candidate tree per plot. The target tree remained the same for all treatment years (2017, 2018 and 2019) but a different tree was measured during the pre-treatment period in 2015. This change was because the plot infrastructure, which determined the CAS system, was not constructed until 2016.

When reporting treatment effects from the present study, we report the *mean enhancement* or *treatment effect*:

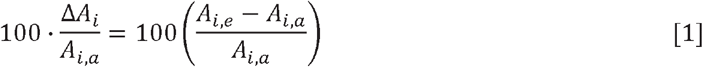

where *A_i,x_* is a measure of gas exchange (*i* = ‘net’ or ‘sat’, see below) at ambient (a) or elevated (e) CO_2_ mixing ratios. When comparing our results with other studies using different eCO_2_ treatments, we report the sensitivity to eCO_2_, following Keenan et al. (2016):

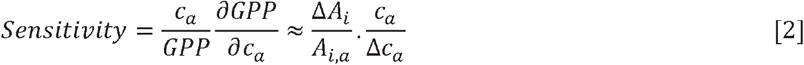

where *c_a_* is the ambient CO_2_ mixing ratio and Δ*c_a_* is the treatment size (e.g. +150 μmol mol^-1^ as in our case). For the conditions of the present study (see ‘Diurnal measurements’ section, below), *c_a_*/Δ*c_a_* = 392/150 = 2.61, and we use net photosynthesis instead of GPP. Hence, our theoretical predicted photosynthetic enhancement (Nowak et al., 2004; Supplemental Appendix 1.) for the + 150 μmol mol^-1^ increase in CO_2_ (i.e., ≈37%;; Hart et al. (2020)), is equivalent to expecting a sensitivity to eCO_2_ of unity.

### Diurnal measurements

Near the canopy top, *in situ* diurnal measurements of gas exchange were conducted on upper canopy oak leaves on 11 and 12 separate summer days of 2018 and 2019, respectively. Measurements of gas exchange (e.g. net CO_2_ photosynthetic assimilation rates, A_net_) were made using a Li-6800 equipped with the default clear Propafilm (Innovia Films Inc., Atlanta, GA) window chamber head, which allowed for natural sunlight to illuminate the leaf. Measurements were conducted in one pair of plots (i.e. one eCO_2_ plot and its paired aCO_2_ plot) on each sampling day. Therefore, each full campaign (n=3) took three days to complete, with the exception of September 2018 and June 2019 where only two replicate plots could be measured. A total of four diurnal campaigns were conducted in both 2018 and 2019, providing a total of 3,426 data points. Five to six healthy leaves were randomly selected in the same oak tree per plot, every 30-40 minutes across the time course of the day for gas exchange measurements, swapping between aCO_2_ and eCO_2_ plots. Measurements were made at the respective growth CO_2_ of aCO_2_ (~405 μmol mol^-1^) or +150 μmol mol^-1^ aCO_2_ (~555 μmol mol^-1^) for eCO_2_ plots, along with other environmental variables such as relative humidity (RH); air temperature (T_air_); and quanta of photosynthetically active radiation (PAR). Measurements were confined to the youngest fully expanded leaves of the leader branch within reaching distance of the CAS system. Measurements were confined to the first flush of leaves across the season for consistency in leaf age. Expanding leaves, judged from colour and texture, were avoided for measurements, as they had not matured in terms of chlorophyll and formation of the photosynthetic apparatus. Once a leaf was inside the chamber, the Li-6800 head was gently positioned and held constant at an angle towards the sun. This was to ensure sun exposure on the leaf, to minimize shading of the chamber head on the measured leaf and to reduce variation across the leaf measurements. Measurements were recorded after an initial stabilisation period (typically ~40 seconds to 1 minute), to meet programmed stability parameters. This allowed for instantaneous steady-state photosynthesis to be captured, yet avoided chamber-related increases in leaf temperature (Parsons et al., 1998). Care was taken to ensure conditions matched those outside the chamber before each measurement was taken. The daily mean RH inside the leaf chamber was between 50% and 77% for all measurements. The mean C_a_ values in the LiCOR chamber head were 390 ± 0.9 μmol mol^-1^ and 538 ± 2.7 μmol mol^-1^, in 2018, and 393 ± 1.0 μmol mol^-1^ and 545 ± 4.8 μmol mol^-1^, in 2019, for aCO_2_ and eCO_2_ respectively. The mean CO_2_ treatments were, therefore, +148 ± 2.8 μmol mol^-1^ in 2018, and +152 ± 4.9 in 2019, and were not statistically different. The gas exchange systems were calibrated before each growing season.

### A-C_i_ curves

*A-C*_i_ curves were conducted in three growing seasons: pre-treatment year (2015), in the first year of CO_2_ fumigation (2017) and third year of CO_2_ fumigation (2019). Measurements were either conducted on attached branches *in situ* (2015 and 2019) or on detached branches harvested by climbers (2017) using a portable open gas exchange system that incorporated a controlled environment leaf chamber (Li-6400XT and LI-6800, LICOR, Inc., Lincoln, NE, USA). Detached branches were transferred to researchers on the ground immediately after excision, where they were placed in a bucket of water to minimize desiccation. Branches were re-cut under water and allowed to stabilize, before starting measurements. Measurement on detached branches were conducted no longer than 45 minutes after collection. Previous studies investigating measurements of gas exchange on severed or attached branches found no significant differences between the two methods (Bader et al., 2016; Verryckt et al., 2020). *A–C*_i_ curves were measured at a *Q* of 1800 μmol m^-2^ s^-1^ (in 2015 and 2019) or 1200 μmol m^-2^ s^-1^ (in 2017) and at a leaf temperature of 25 °C. Before each curve, a stabilisation period of between 5 to 10 minutes was used depending on the prevailing environmental conditions and each curve took an average of 40 minutes. Light-saturated net photosynthesis (A_sat_) were estimated from *A–C*_i_, curves at growth [CO_2_]. The CO_2_ concentrations were changed in 12 to 14 steps starting at the respective growth [CO_2_]; every 100 μmol mol^-1^ down to 50 μmol mol^-1^ (near the photosynthetic CO_2_ compensation point), then increasing to 1800 μmol mol^-1^ in roughly 200 μmol mol^-1^ increment steps. Five to six replicate *A–C*_i_ curves on different leaves per CO_2_ treatment were measured per day. Measurements were taken between 09:00 and 11:00, and 14:00 and 17:00 to avoid potential midday stomatal closure (Valentini et al., 1995). Measurements were made using the treatment pair arrangement of one aCO_2_ and one eCO_2_ plot per day (n=3).

### Leaf carbon and nitrogen

Oak leaves were collected from the top of the canopy in each month, May to November in 2015 and 2019, by arborist climbers, and stored immediately at −25 °C. Two upper canopy leaves, from one tree per plot, were selected for elemental analyses, these trees corresponded to the measurement tree for leaf gas exchange. Each leaf was photographed on white graph paper, with a ruler for reference. Leaf area analysis was conducted using imaging software Image J (IMAGE J v1.53, National Institutes of Health, Bethesda, MD, USA) and the fresh weight was recorded. Each leaf was oven dried at 70 °C for at least 72 hours, re-weighed for dry weight and the leaf mass per unit area was calculated. Dried leaf fragments were ground and each sample (~2 mg) was enclosed in a tin capsule. Samples were analysed for *δ*^13^C, total C, and total N using an elemental analyser interfaced with an isotope ratio mass spectrometer (Sercon Ltd., Cheshire, UK).

### Statistical analysis

All statistical analysis were performed in R version 4.0.3 (R Core Team, 2020). Before statistical analysis, all data were checked for normality by inspection of the Q–Q. plots and Levene’s test, and residuals from model fitting were checked for evidence of heteroscedasticity. Hourly averages of diurnal measurements were analysed using a linear mixed effects model (‘*lmer*’ package). Fixed categorical factors in this model were CO_2_ treatment (i.e., aCO_2_ or eCO_2_), sampling month and sampling year (i.e. 2018 or 2019), in addition to their interactions. Additionally, ‘time of day’ and ‘plot’ were represented as random factors, the latter as individual trees were nested within each experimental plot. Type III F-statistics associated with the mixed model analysis (repeated-measures analysis of variance (ANOVA)) were reported. Statistically significant CO_2_ treatment differences among groups were further tested with Tukey’s post hoc test using the R package ‘emmeans’ (*P* < 0.05 reported as significant). To investigate the dependence of photosynthetic enhancement with variation of light, the diurnal gas exchange data, with leaf temperature, T_leaf_ > 18 °C, and vapour pressure deficit (*D*), *D* < 2.2 kPa, were sub-divided into four light (*Q*) categories, each sampled about equally. The *Q* classes were chosen based on the characteristic response of A_net_ to light as follows: *Q* < 250; 250 ≤ *Q* < 500; 500 ≤ *Q* < 1000; and *Q* ≥ 1000 μmol m^-2^ s^-1^. CO_2_ treatment, year, and *Q* category were then used as parameters in the ANOVA.

The photosynthetic CO_2_ response (*A–C*_i_) curves were fit with the model of Farquhar et al. (1980) to estimate the apparent maximum rate of photosynthetic Rubisco carboxylation (*V*_cmax_, μmol m^-2^ s^-1^) and the apparent maximum rate of photosynthetic electron transport (*J*_max_, μmol m^-2^ s^-1^) using *‘Plantecophys’* package in R (Duursma, 2015). The model-fitting was undertaken to provide insight into photosynthetic capacity and its response to long-term exposure to elevated [CO_2_] (Rogers & Ellsworth, 2002). We tested for outliers by examining the *J*_max_/*V*_cmax_ ratio, RMSE values and standard errors (SE) for fits of *J*_max_ and *V*_cmax_, all of which indicate violations to the theory for fitting these curves (Sharkey et al., 2007). Visual inspection of each *A-C*_i_ curve with outliers allowed us to identify any incomplete curves and/or mechanical failures and those curves were subsequently removed. This accounted for < 10% of the data, leaving a total of 86 *A–C*_i_ curves across the three sampling years in the analysis.

## Results

### Measurement conditions

Overall, diurnal measurements were conducted on dry, sunny days (Fig. 1), and environmental conditions (*Q* and T_leaf_) were consistent between aCO_2_ and eCO_2_ across the two growing seasons of diurnal measurements (Figs. 2A, B and 3A, B). *Q* levels were largely comparable between CO_2_ treatments although cloud and temperature conditions were more variable among sampling days and campaigns in 2018 than in 2019.

**Figure 2.**
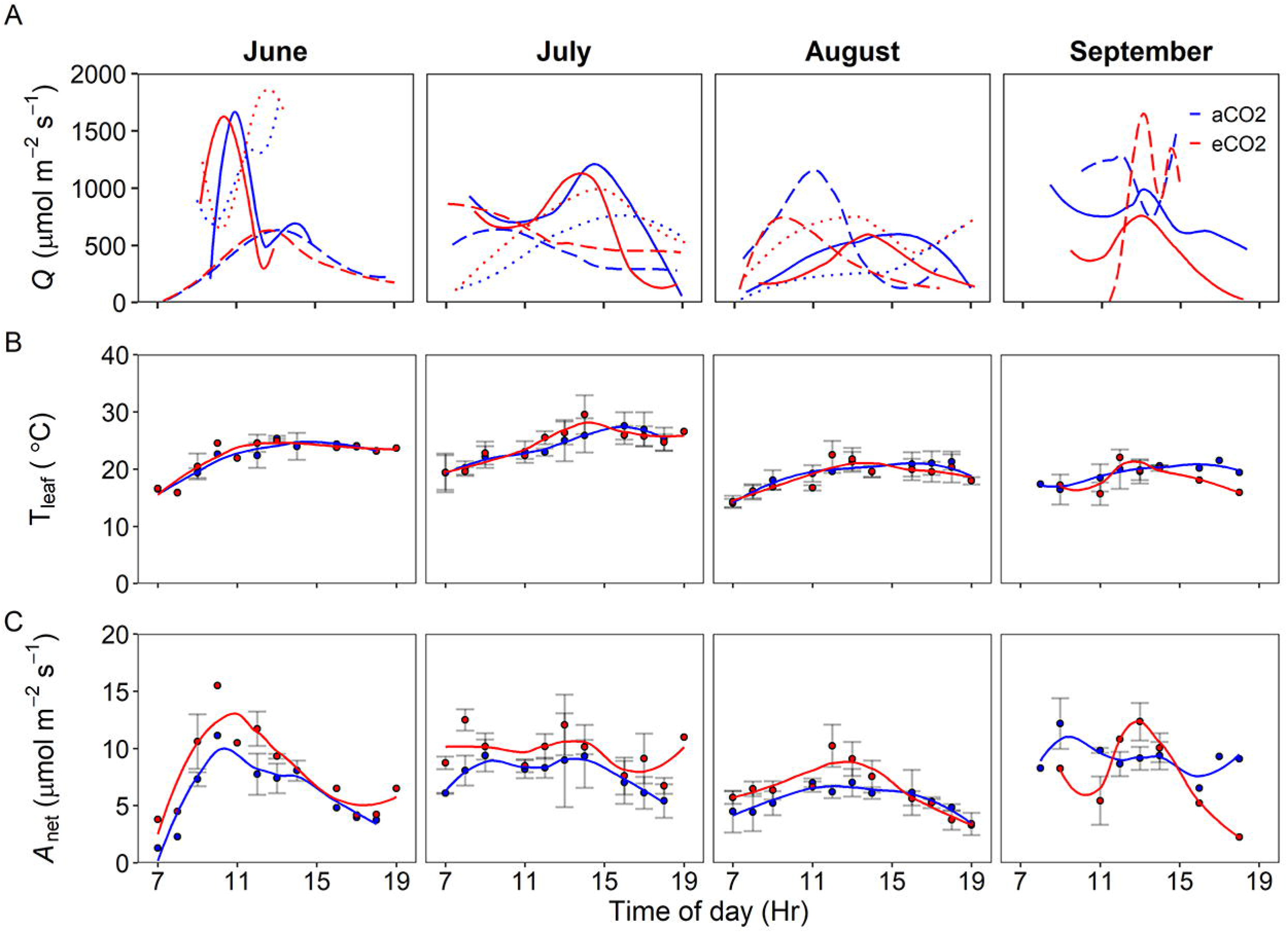
*In situ* diurnal measurements of **A)** *Q* (μmol m^-2^ s^-1^), **B)** hourly mean T_leaf_ (°C) and **C)** hourly mean A_net_ (μmol m^-2^ s^-1^); each fitted with an LOESS regression, at BIFoR FACE in 2018 from the upper *Q. robur* canopy. Error bars indicate n=3, with the exception of September where only two replicate plots were measured and not all time points were replicated. The line types in A) represent replicate plot pairings of; plots 1 and 3 (dotted), plots 2 and 4 (solid), and plots 5 and 6 (long-dash) and the two colours represent the CO_2_ treatments of aCO_2_ (blue) and eCO_2_ (red).

**Figure 3.**
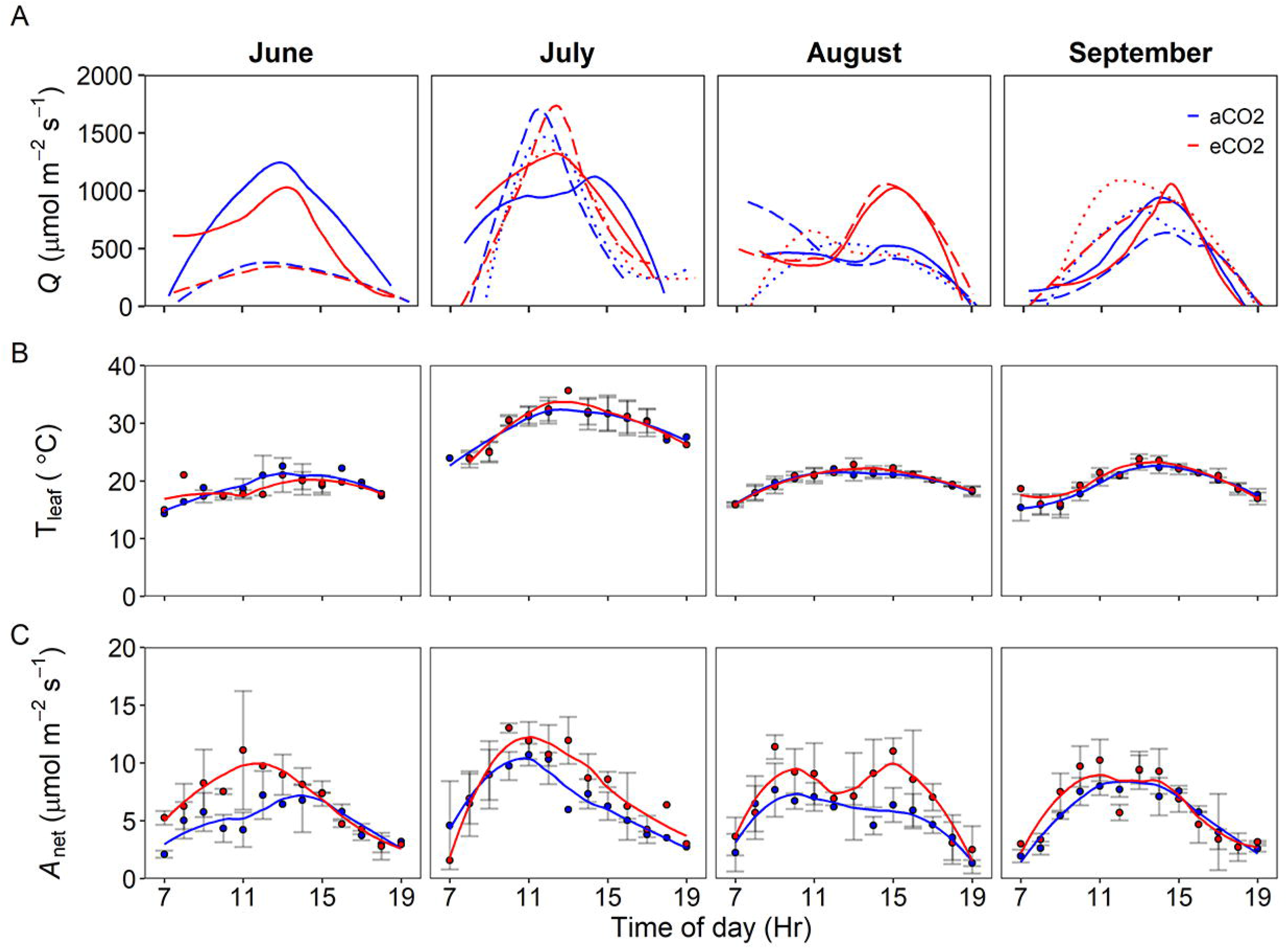
*In situ* diurnal measurements of **A)** *Q* (μmol m^-2^ s^-1^), **B)** hourly mean T_leaf_ (°C) and **C)** hourly mean A_net_ (μmol m ^2^s^-^); each fitted with an LOESS regression, at BIFoR FACE in 2019 from the upper *Q. robur* canopy. Error bars indicate n=3, the exception of June where only two replicate plots were measured and not all time points were replicated. The line types in A) represent replicate plot pairings of; plots 1 and 3 (dotted), plots 2 and 4 (solid), and plots 5 and 6 (long-dash) and the two colours represent the CO_2_ treatments of aCO_2_ (blue) and eCO_2_ (red).

Leaf temperature was more stable than *Q* with lower variability across the diurnal sampling, high similarity between sampling days, and high consistency between CO_2_ treatments. There were differences of up to 15 °C in midday measurements of T_leaf_, between months, suggesting a seasonal influence as would be expected from the site’s mid-latitude location, with differences more prominent in 2019 than 2018. The highest T_leaf_ values were observed in July with a common seasonal decline after this campaign.

Analysis of the diurnal dataset showed the range of mean daily A_net_ was similar between years, however the highest mean daily A_net_ (12.2 μmol m^-2^ s^-1^) was reported in 2018. Contrasting seasonal patterns were observed between the sampling years of 2018 and 2019, with decreases in mean daily A_net_ across the growing season observed in 2018 compared to increases in A_net_ in 2019. In both sampling years, we observed a significant enhancement of A_net_ when exposed to eCO_2_ (P <0.05, Table 1 and Figs. 2 and 3). Here, we did not observe any significant effect of either season or sampling year on A_net_ (Table 1.). Therefore, from measurements of A_net_ collected from the diurnal dataset, a mean eCO_2_-driven photosynthetic enhancement (i.e., 100. Δ*A_i_/A_i,a_*) of 23 ± 4% was observed across the 2-year period of this study.

**Table 1.**
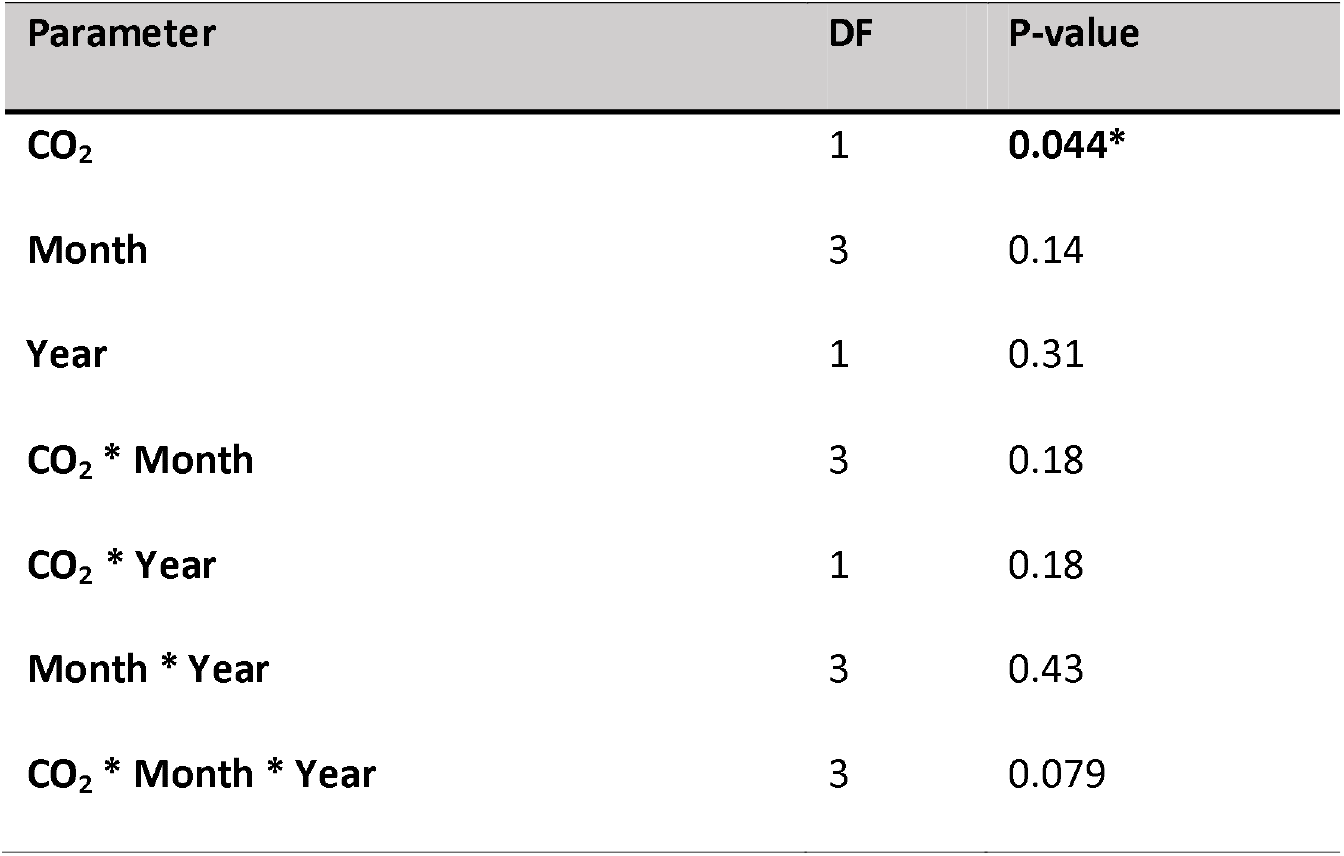
Linear mixed-effects model analysis for photosynthesis with CO_2_ treatment (CO_2_) using the diurnal dataset, sampling month (Month) and sampling year (Year) as fixed factors and random effects of ‘plot’ and ‘time’. Type III sums of squares computed using restricted maximum likelihood estimates for *F*-tests. The numerator degrees of freedom (df) for each *F*-test are shown. A post-hoc Tukey test was used to determine the significance relationships. Significance of CO_2_ treatment is noted in the rightmost column as (* = P < 0.05).

### Photosynthesis and variation in photon flux density (Q)

This study analysed the role of measurement *Q* affecting A_net_ and its response to eCO_2_ in separate growing seasons to investigate photosynthetic enhancement values at different light conditions. In each light category (see Methods, above), the light conditions between the CO_2_ treatments were statistically comparable (Figure 4, Supplementary table 1: S1). Mean, median, and interquartile range of A_net_ increased with increasing *Q* class for both sampling years and CO_2_ treatments (Fig. 4A, Table 2). We observed no significant effect of year for A_net_ in this study, but we did observe a larger variation in A_net_ in 2019, when compared to 2018 (Table 2, Fig. 4A). Values of mean A_net_ ranged from 4.6 ± 0.3 μmol m^-2^ s^-1^, at the lowest *Q* level with a mean of 150 μmol m^-2^ s^-1^, to 11.5 ± 0.7 μmol m^-2^ s^-1^ at highest *Q* (mean *Q* of 1360 μmol m^-2^ s^-1^). Additionally, in both sampling years A_net_ was significantly higher under eCO_2_ conditions when compared to aCO_2_ (P < 0.05, Table 2, Fig. 4A). Consistent with our hypothesis, we observed mean eCO_2_-driven photosynthetic enhancement to increase with increasing *Q*, with the largest enhancement observed at highest *Q* in both sampling years, 30 ± 9%, and 35 ± 13%, for 2018 and 2019 respectively (Fig. 4B). In 2018, eCO_2_-driven photosynthetic enhancement ranged from 7 ± 10%, in the lowest *Q* class, to 30 ± 9%, in the highest *Q* class (Fig. 4B). A similar positive relationship between eCO_2_-driven photosynthetic enhancement and *Q* was present in 2019 with enhancement ranging from 11 ± 6%, in the lowest *Q* class, to 35 ± 13%, in the highest *Q* class (Fig. 4, B). There was no significant effect of year (Table 2) and therefore the mean eCO_2_-driven photosynthetic enhancement at light saturation (i.e. in the highest *Q* class) was on average 33 ± 8 % across the two sampling years. Our results report that the mean eCO_2_-driven photosynthetic enhancement of light-saturated A_net_ (A_sat_) in both sampling years was consistent, within error (using 95 % confidence intervals), of the theoretical predicted enhancement based on proportion of CO_2_ increase (≈37 ± 6%), indicating a sensitivity to eCO_2_ (equation 2, above) of close to unity for A_sat_.

**Figure 4.**
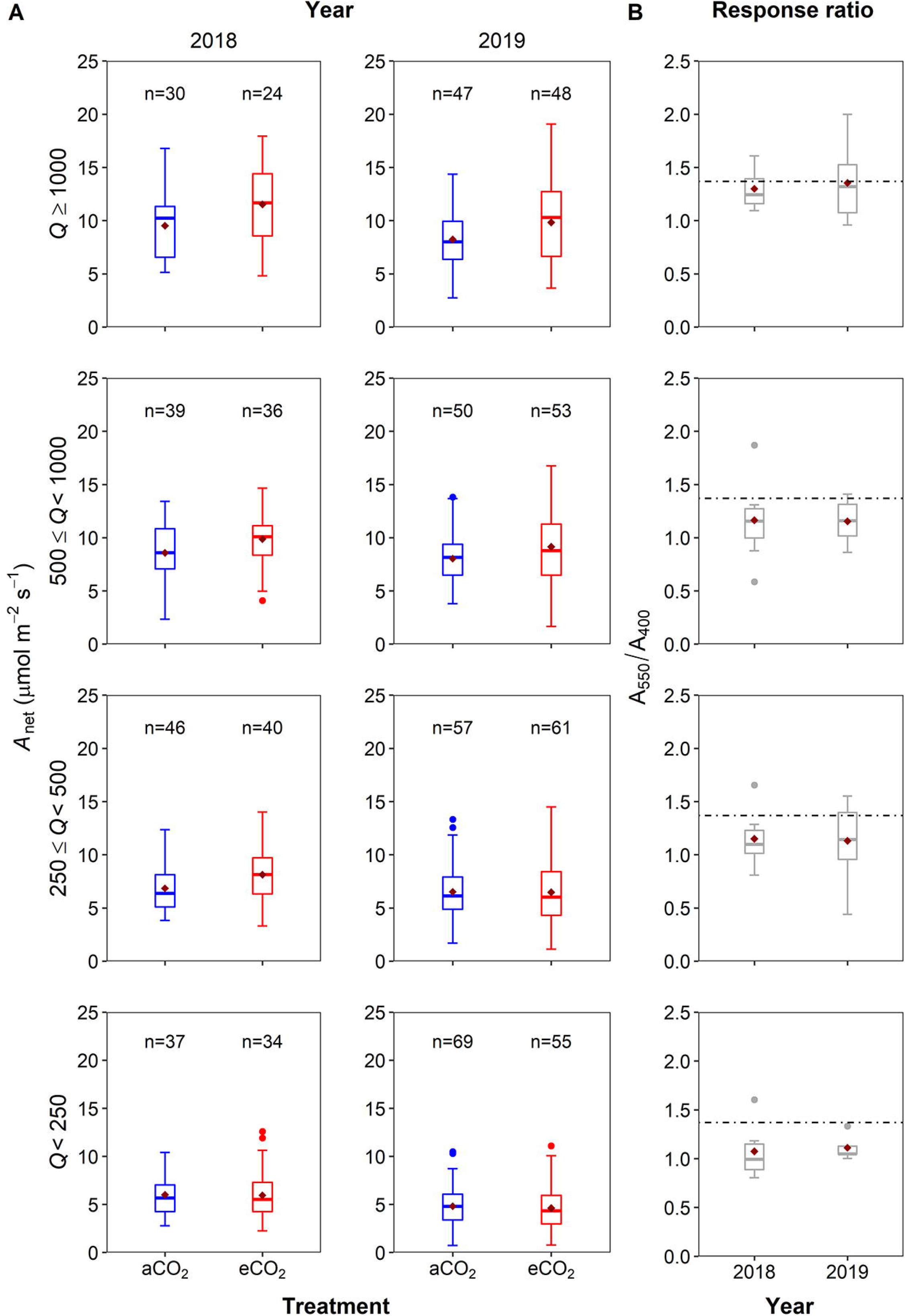
A) The distribution of net photosynthesis (A_net_) (μmol m^-2^ s^-1^) in each of the four photon flux density (*Q*) categories (*Q* < 250; 250 ≥ *Q* < 500; 500 ≥ *Q* < 1000; and *Q* ≥ 1000 μmol m^-2^ s^-1^) for years 2018 (left) and 2019 (right). Whiskers denote the 5 %ile and 95 %ile; outliers are plotted as individual points (filled circles). The box denotes the interquartile range and the bar denotes the median with the number of data points above each boxplot. The mean is also plotted as a diamond symbol. Data uses diurnal gas exchange measurements in the upper canopy oak trees at the BIFoR FACE facility with T_leaf_ >18 °C and *D* <2.2 kPa, in eCO_2_ (red) or aCO_2_ (blue) treatments. Red diamonds indicate the mean A_net_ values. B) Boxplots of the enhancement response ratio (A_550_/A_400_) (grey) for each year, and predicted enhancement ratio (dashed line) (1.37) following Nowak et al (2004).

**Table 2.** Linear mixed-effects model parameters for prediction of A_net_ with variation in photo flux density (*Q*). Type III sums of squares computed using restricted maximum likelihood estimates for *F*-tests. The numerator degrees of freedom (DF) for each *F*-test are shown. A post-hoc Tukey’s test was used to determine the significance relationships. Significance is noted in the rightmost column as (*\*\*\* = P* <0.001; ** = *P* < 0.01; *=*P* < 0.05)

| Parameter | DF | P-value |
| --- | --- | --- |
| **CO<sub>2</sub>** | 1 | **0.016\*** |
| **Year** | 1 | 0.062 |
| **Q** | 3 | **<0.001\*\*\*** |
| **CO<sub>2</sub> \* Year** | 1 | 0.97 |
| **CO<sub>2</sub> \* Q** | 3 | **0.011\*** |
| **Year \* Q** | 3 | **0.0078\*\*** |
| **CO<sub>2</sub> \* Year \* Q** | 3 | 0.13 |

### Photosynthetic capacity and foliar nitrogen

The seasonal and interannual biochemical changes in *Q. robur* were assessed via differences in leaf apparent maximum CO_2_ carboxylation capacity (*V*_cmax_) and apparent maximum electron transport capacity for RuBP regeneration (*J*_max_) (Fig. 5.) to assess the photosynthetic capacity in the initial years of the long-term experiment. Initially, we tested for differences between the year of sampling and found no statistical difference of either *V*_cmax_ or *J*_max_ between the three sampling years (2015, 2017 and 2019) (Fig. 5, Supplementary table 2: S2). This study found no significant effects of CO_2_ enrichment on *V*_cmax_ or *J*_max_ across the two years of CO_2_ enrichment, i.e. the 1^st^ and 3^rd^ years, and no significant effect of season between the three measurement years (Fig. 5, Table 3.). However, this study did observe a significant effect of month for the variable *V*_cmax_ in 2019, whereby an increase in *V*_cmax_ was observed with progression of the growing season (Fig. 5A, Table 3.). Thus, this study observed no statistical evidence to suggest photosynthetic down-regulation of either *V*_cmax_ or *J*_max_ under elevated CO_2_ across the three years of eCO_2_ exposure in *Q. robur*.

**Figure 5.**
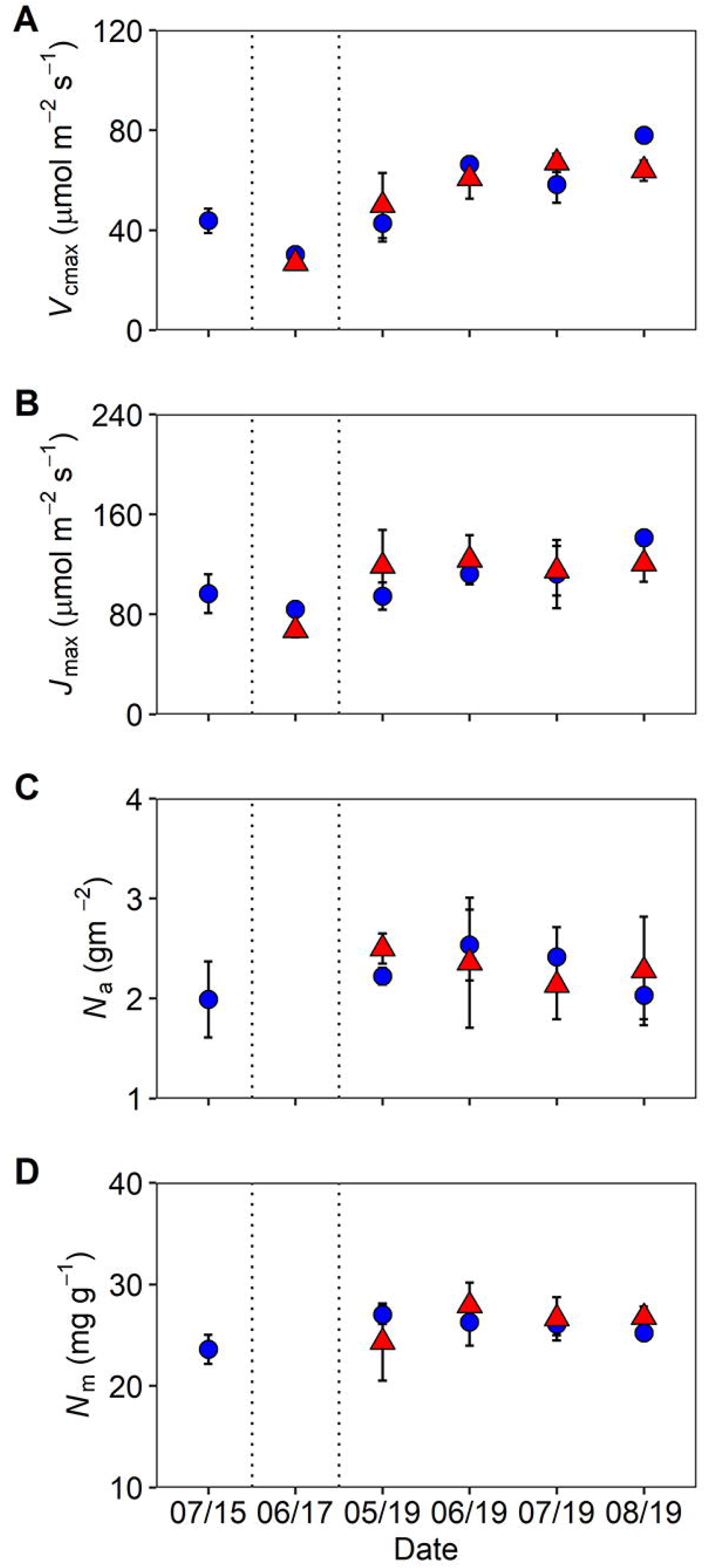
Maximum rates of (A) carboxylation (*V*_cmax_) and (B) electron transport (*J*_max_), in addition to (C) area based (*N_a_*) and D) mass based (N_*m*_) leaf nitrogen of upper canopy *Q. rohur* from 2015 to 2019 at BIFoR FACE. Means (± SE) of whole-plot averages (n=3) for ambient (blue circles) and elevated (red triangles) CO_2_ treatments. Dashed line indicate the separation of sampling years with campaigns labelled ‘month/year’, as follows: Pre-treatment (‘07/15’); 1^st^ Year (‘06/17’); and the 3^rd^ year (‘05/19’ - ‘08/19’) of CO_2_ fumigation. Data points may obscure error bars.

**Table 3.** Linear mixed-effects model analysis for *V*_cmax_, *J*_max_, net photosynthesis (A_net_), area-based leaf nitrogen (N_*a*_) and mass-based leaf nitrogen (N_*m*_) with CO_2_ treatment (CO_2_) and sampling month (Month) as fixed factors and random effects of ‘plot’ and ‘time’. Type III sums of squares computed using restricted maximum likelihood estimates for *F*-tests. The numerator degrees of freedom (DF) for each *F*-test are shown. Significance is noted in boldface as (* *P* < 0.05)

| Parameter | | $V_{\text{cmax}}$ | $J_{\text{max}}$ | $A_{\text{net}}$ | $N_m$ | $N_a$ |
| --- | --- | --- | --- | --- | --- | --- |
| | DF | $P$- value | $P$- value | $P$- value | $P$- value | $P$- value |
| $\text{CO}_2$ | 1 | 0.70 | 0.37 | **0.042\*** | 0.42 | 0.64 |
| Month | 3 | **0.02\*** | 0.15 | **0.034\*** | 0.93 | 0.052 |
| $\text{CO}_2$ * Month | 3 | 0.20 | 0.57 | 0.33 | 0.69 | 0.11 |

Consistent with previous research, this study observed a strong positive linear relationship between *J*_max_ and *V*_cmax_, which remained unchanged across CO_2_ treatments and growing season (r^2^ = 0.75 ambient; r^2^ = 0.71 elevated) (Supplementary Figure 3.). Additionally, no eCO_2_-induced decreases in either area-based foliar nitrogen (N_a_) or mass-based foliar nitrogen (N_m_) were observed (Fig. 5; C and D, Table 3.) across the study period. No change in foliar nitrogen is corroborative of the results in Fig. 5 and also suggest the absence of photosynthetic downregulation under eCO_2_ in mature *Q. robur* in the first three years of the long-term experiment.

The instantaneous response ratio (2015) and the longer-term response ratio (2017 and 2019) were calculated using the light-saturated A_net_(i.e., A_sat_) values at growth CO_2_ from the *A–C*_i_ datasets (Fig. 6B). There was no significant difference between the measurement year in either A_sat_ or the response ratio suggesting comparability between the instantaneous response ratio and the longer-term response ratio (Supplementary table 3: S3). A significant treatment effect was observed for A_sat_ (Fig. 6A, Table 3.) in all three sampling years, with a mean eCO_2_-driven photosynthetic enhancement of 24 ±2%, 31 ± 7% and 32 ± 11% in 2015, 2017 and 2019, respectively, under elevated CO_2_ when compared to aCO_2_. A significant effect of month on A_sat_ was observed in 2019, with A_sat_ increasing with the progression of the growing season (Table 3, Fig. 6A). The photosynthetic enhancement observed from our *A–C*_i_ curve datasets are consistent with the values obtained in the diurnal dataset (33 ± 8%, Fig. 5) but is lower than the theoretical predicted enhancement calculated via CO_2_ increase (37%) (Supplemental Appendix 1.). In summary, the consistency in the two separate measurements (i.e. diurnal and *A–C*_i_ curves) support the finding of sustained eCO_2_-driven photosynthetic enhancement in mature *Q. robur* across the first three years of the BIFoR FACE experiment.

**Figure 6.**
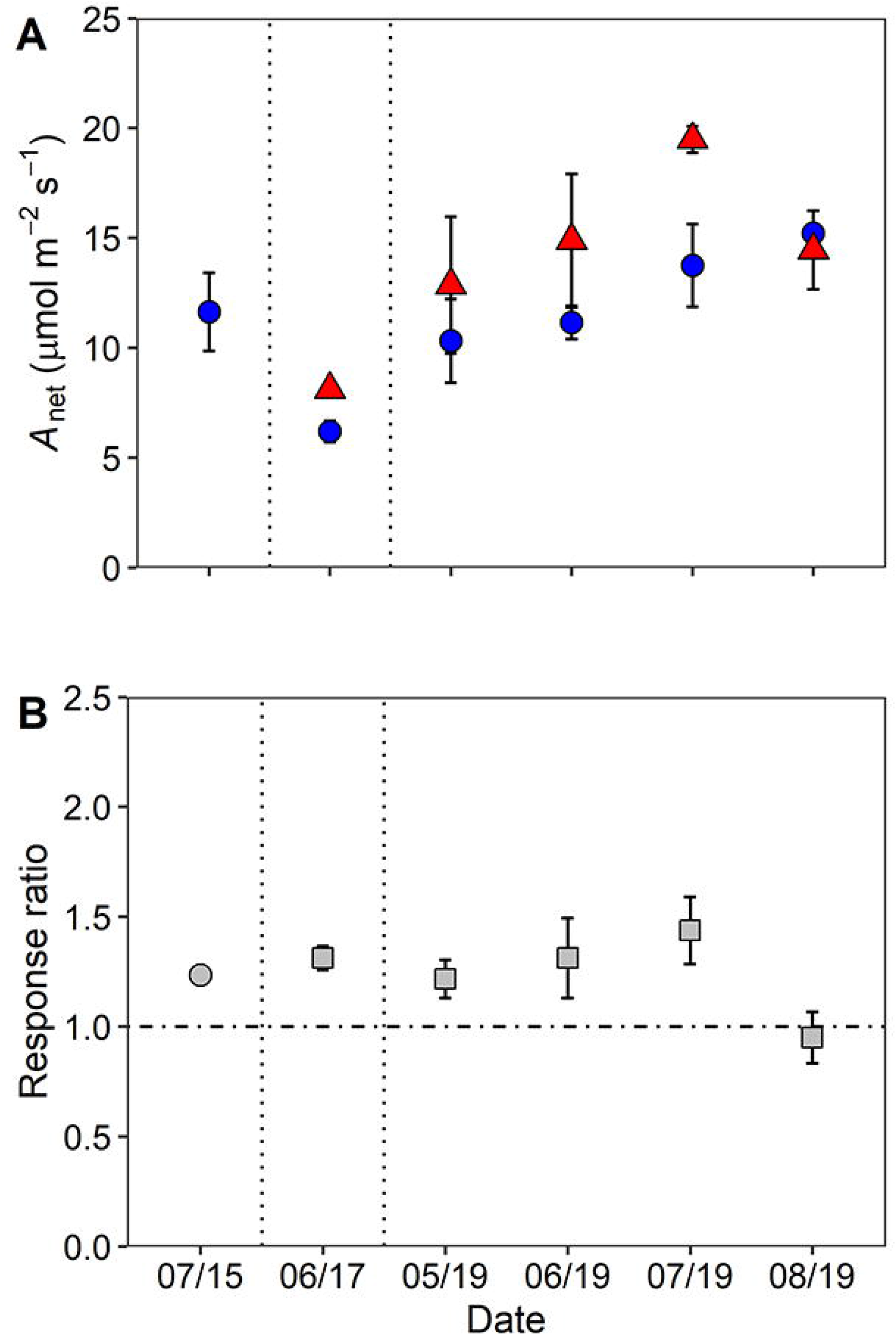
A) Net photosynthesis (A_net_) at growth CO_2_ and B) Instantaneous (2015) and longer-term (2017 and 2019) response ratios in the upper oak canopy using the *A–C*_i_, curve data. Means (± SD) of the plots per treatment are shown across six sampling campaigns for aCO_2_ (blue circles), eCO_2_ (red triangles) and either the instantaneous (grey squares) or longer-term response ratio (grey circles). Dashed line indicate the separation of sampling years with campaigns labelled as follows; Pre-treatment (‘07/15’), 1^st^ Year (‘06/17’) and the 3^rd^ year (‘05/19’-‘08/19’) of CO_2_ fumigation.

## Discussion

There is ample data on the short-term enhancement of photosynthesis by eCO_2_ in young trees using a variety of experimental set-ups from tree chambers to FACE experiments (e.g. Ainsworth & Rogers, 2007, Crous et al. 2013), but few data for mature forest-grown trees with multi-year CO_2_ exposure in a FACE setting. For mature trees, available evidence suggests that there are significant increases in light-saturated A_net_ (Ellsworth et al., 2017; Körner et al., 2005) but there have been mixed results regarding the magnitude of photosynthetic enhancement (range 13-49% per 100 ppm of CO_2_ increase) and occurrence of photosynthetic downregulation in mature forest-grown trees (Bader et al., 2010, 2016; Crous et al., 2008; Ellsworth et al., 2017; Warren et al., 2015). In this study, we predicted a theoretical A_net_ enhancement of 37% for the 150 μmol mol^1^ increase in CO_2_ at BIFoR FACE following reasoning in Nowak et al. (2004) (Supplemental Appendix 1). After three years of eCO_2_ exposure in mature temperate oak forest, net photosynthetic rates of upper canopy foliage from *Q. robur* were on average 23 ± 4% higher, based on the diurnal dataset, in the trees exposed to elevated CO_2_ when compared to control plots (Figs. 2, 3 and 4; Tables 1 and 3). The eCO_2_-driven photosynthetic enhancement observed is substantially lower than the theoretical expected enhancement of 37%, likely due to diurnal and seasonal variation in prevailing environmental conditions such as lower air temperatures, lower light conditions, and varying vapour pressure deficits. Only considering light-saturated A_net_ (A_sat_) from the diurnal dataset, our mean photosynthetic enhancement is greater than the average diurnal enhancement, at 33 ± 8% rather than 23%. Furthermore, our independent estimate of A_sat_ enhancement based on the *A–C*_i_ curve data is 32 ± 11%, which is comparable within error (using 95% confidence intervals) to both the A_sat_ value from the diurnal measurements and the hypothesized enhancement of 37%. A slight stomatal closure in eCO_2_ could have contributed to the slightly lower photosynthetic enhancement than the hypothesized enhancement of 37% (See Supplemental Appendix 1). However, our average light-saturated photosynthetic enhancement is generally lower than previously reported values in canopy dominant trees from other forest FACE experiments (Bader et al., 2010, 42-48%; Crous et al., 2008, 40-68%; Liberloo et al., 2007, 49%; Sholtis et al., 2004, 44%), but is somewhat higher than the value of 19% from the EucFACE experiment on mature *Eucalyptus* trees (Ellsworth et al. 2017). The lower photosynthetic enhancement observed at EucFACE was likely due to lower nutrient availability compared to BIFoR (Crous et al., 2015), although there were other differences such as the tree species and prevailing temperatures that would also affect the magnitude of the photosynthetic enhancement.

### The role of environmental conditions for photosynthetic enhancement

Consistent with our initial hypothesis, we observed significantly higher A_net_ and a 24% higher photosynthetic enhancement under the highest light conditions at BIFoR FACE (i.e., *Q* > 1000 μmol m^-2^ s^-1^) compared to the lowest light category. Thus, a negative linear relationship was observed for both A_net_ and eCO_2_-induced photosynthetic enhancement with decreasing light levels. Our results are consistent with previous research on mature trees that observed an effect of light on the magnitude of CO_2_-driven stimulation of photosynthesis (Bader et al., 2016), suggesting variation in light should be considered when assessing the response to eCO_2_.Consequently, the relationship of A_net_ and CO_2_ treatment effect with light intensity is important when scaling upper canopy data both across diurnal periods of light limitation and extending to the whole canopy, of shaded and sunlit leaves, to avoid overestimating canopy-scale photosynthesis by temperate forests.

It has been previously suggested that larger photosynthetic enhancement may be expected in low light environments (Hättenschwiler, 2001; Norby & Zak, 2011). For example, deep shaded tree seedlings displayed greater photosynthetic gains than those in moderate shade (photosynthetic enhancement of 97% and 47%, respectfully) with exposure to eCO_2_ (Kitao et al., 2015). In light-limited environments, higher CO_2_ concentrations can increase the apparent quantum yield and reduce the light compensation point (LCP) leading to enhanced carbon uptake (Larcher, 2003; Kitao et al., 2015). Hättenschwiler (2001) found large interspecific variability and, in *Quercus*, that greater photosynthetic responses to CO_2_ occurred under higher light when compared to low light. However, both Kitao et al., (2015) and Hättenschwiler (2001) studied tree seedlings in contrast to upper canopy leaves of a canopy dominant species in the present study. Although shade leaves were not measured here, the results here from the top of the tree canopy provide an important benchmark for the magnitude of photosynthetic enhancement by elevated CO_2_ in a mature oak forest.

In addition to light intensity, the photosynthetic response of *Q. robur* varied across the growing season, as has been observed in many other trees (Rogers & Ellsworth, 2002; Sholtis et al., 2004; Tissue et al., 1999). Here, A_sat_ (derived from the *A–C*_i_ dataset) in both CO_2_ treatments increased about 50% from early in the season (May), to the middle of the season (July); yet, the relative response ratio to eCO_2_ was stable throughout this period at 32%. Additionally, when assessing the diurnal dataset, we found contrasting seasonal patterns between 2018 and 2019, with decreases in A_net_ across the growing season observed in 2018 compared to increases in A_net_ in 2019, likely due to drier and warmer conditions in 2018. Previous research has identified reductions in photosynthesis across the season is largely associated with drier conditions (Gunderson et al., 2002), which support the results observed in the present study. This suggests that the influence of soil water availability on the seasonal pattern in oak physiology is critical for determining seasonal C-uptake by mature forests and should be further investigated in mature *Q. robur* to improve longer term carbon-climate models (see Limousin et al., 2013).

Previous research has identified eCO_2_-driven photosynthetic responses observed in seedlings and saplings may not reflect the photosynthetic responses of mature forest-grown trees (Hättenschwiler et al., 1997). The present study provided a unique opportunity to assess the eCO_2_-driven photosynthetic responses in 175-year old canopy dominant trees and found lower photosynthetic stimulation than the many previous studies on tree seedlings and younger trees (e.g. Ainsworth & Long, 2005; Curtis & Wang, 1998; Crous et al., 2008; Liberloo et al., 2007; Sholtis et al., 2004). The age dependency of CO_2_ responsiveness to photosynthesis in trees (Turnbull et al., 1998; Wujeska-Klause et al., 2019a), highlights the importance of long-term experiments, such as the present study and others in understanding potential variable responses across the lifetime of a tree, vital for accurate climate-carbon modelling of forests.

### Did changes to photosynthetic capacity or leaf biochemistry occur under eCO_2_?

In some studies, a time-dependent decline in the magnitude of eCO_2_-induced photosynthetic enhancement, i.e. photosynthetic downregulation, has been observed (Cure & Acock, 1986; Gunderson & Wullschleger, 1994). Here, we hypothesized that there may be reductions in *V*_cmax_, *J*_max_ and leaf N, particularly in the 3^rd^ year of eCO_2_ exposure (Luo et al., 2004). Our analysis of the 86 *A–C*_i_ curves collected in this experiment revealed no decrease in the rate of *V*_cmax_ or *J*_max_, indicating that there were no significant changes in the photosynthetic capacity of *Q. robur* over the first three years of exposure to elevated CO_2_. A lack of photosynthetic downregulation has also been found in similar seasonally deciduous species, including the closely related species *Quercus petraea* (Bader et al., 2010), in addition to *Liquidambar styraciflua, Populus spp*. and *Betula papyrifera* (Herrick & Thomas, 2001; Sholtis et al., 2004; Liberloo et al., 2007; Uddling et al., 2009). An apparent lack of downregulation has also been observed in other mature forest-grown species (Ellsworth et al., 2017; Bader et al., 2010).

As nitrogen is required for the synthesis and maintenance of photosynthetic proteins, eCO_2_-driven photosynthetic downregulation has been associated with declines in foliar N (as reviewed in Medlyn et al., 1999) and soil N-limitations (e.g. Crous et al., 2008; Rogers & Ellsworth, 2002; Warren et al., 2015). The current study on *Q. robur* did not find any changes in either mass- or area-based leaf nitrogen across the study period, indicating there are no reductions to photosynthetic capacity (Fig. 5.). This corroborates the findings from the *V*_cmax_ and *J*_max_ parameters, supporting the suggestion for sustained photosynthesis in *Q. robur* over the first three years of exposure to elevated CO_2_. Hence, there were no changes to the ratio of *J*_max_ to *V*_cmax_, indicating that the relationship between carboxylation and light-harvesting processes was not affected by CO_2_ treatment, as found in previous studies (Crous et al., 2008; Medlyn et al., 1999), including the closely related species, *Q. petraea* (Bader et al., 2010). These results may point to soil nutrient availability not yet limiting the photosynthetic processes in this forest system. The BIFoR FACE site receives moderately high atmospheric N deposition (~22 Kg N/ha/yr) thought to represent 15% of the total nitrogen nutrition of temperate deciduous trees, likely preventing ecosystem N-limitation at present (Rennenberg & Dannenmann, 2015). Therefore, with adequate N deposition in the soil, sustained photosynthetic enhancement was observed in the first three years of eCO_2_ exposure at BIFoR FACE.

## Conclusions

After three years of eCO_2_ exposure in a temperate deciduous forest at the BIFoR FACE facility, photosynthetic enhancement of mature *Q. robur* leaves at the top of the canopy was sustained across all years and was 33 ± 8% (mean ± s.e.) at light saturation, close to the theoretical expectation. The magnitude of photosynthetic enhancement was significantly affected by light conditions with higher enhancement at higher light. We found no evidence of photosynthetic downregulation under eCO_2_ and no declines in leaf nitrogen in the upper canopy. The lack of evidence for downregulation suggest there are sufficient soil nutrients for *Q. robur* to maintain a relatively high photosynthetic enhancement under eCO_2_ conditions, at least to this point in the eCO_2_ experiment. Much further work remains to determine the movement and allocation of this enhanced C uptake in the forest. Our results are consistent with a sustained, positive C uptake response to rising atmospheric CO_2_ in a mature deciduous forest tree species, provided adequate nutrients are available.

## Acknowledgments

We thank the BIFoR technical team for canopy access operations and Ian Boomer for technical support with leaf elemental analysis. AG gratefully thanks Agnieszka Wujeska-Klause for guidance with statistical analysis in the early stages of the manuscript. AG gratefully acknowledges a studentship provided by the John Horseman Trust and the University of Birmingham. The BIFoR FACE facility is supported by the JABBS foundation, the University of Birmingham and the John Horseman Trust. ARMK acknowledges support from the Natural Research Council through grant (NE/S015833/1) which also facilitated DSE’s participation. We further gratefully acknowledge advice and field measurement collection in the first CO_2_ fumigation season from Michael Tausz and Sabine Tausz-Pösch, respectively.

## Author contributions

ARMK, JP, and AG designed the study; AG, KYC and DSE collected the data. AG organised the datasets under the supervision of DSE, with input from ARMK; AG and DSE designed and performed the statistical analyses, with input from KYC and ARMK. AG and DSE wrote the first draft of the paper. All authors contributed to the manuscript revision, and read and approved the submitted version.

## Symbols and Abbreviations

[CO_2_] CO_2_: concentration of the atmosphere
*A*: photosynthesis
*A-C*_i_: curve Photosynthetic CO_2_ response curve
aCO_2_: CO_2_ at ambient Ca (~405 ppm)
A_net_: photosynthetic rates.
A_sat_: Light-saturated net photosynthesis
C: Carbon
CAS: Canopy access system
*C*_i_: CO_2_ concentration of the intercellular leaf space
eCO_2_: CO_2_ at elevated Ca (+150 ppm ambient)
FACE: Free air carbon dioxide enrichment
*J*_max_: Maximal photosynthetic electron transport rate (a proxy for ribulose-1,5-bisphosphate regeneration)
N: Nitrogen
N_a_: Area-based foliar Nitrogen
N_m_: Mass-based foliar Nitrogen
*Q*: photon flux density
RH: relative humidity
T: temperature
T_air_: Air temperature
T_leaf_: Leaf temperature
SE: Standard error of the mean
V_cmax_: Maximal carboxylation rate of Rubisco
*D*: vapour pressure deficit of the atmosphere
δ^13^C: ratio of ^13^C to ^12^C stable carbon isotopes
δ N ratio: of ^15^N to ^14^N stable carbon isotopes

