## Supplementary material for "Is photosynthetic enhancement sustained through three years of elevated CO_2_ exposure in 175-year old *Quercus robur*?"

**Supplementary Figure 1.**

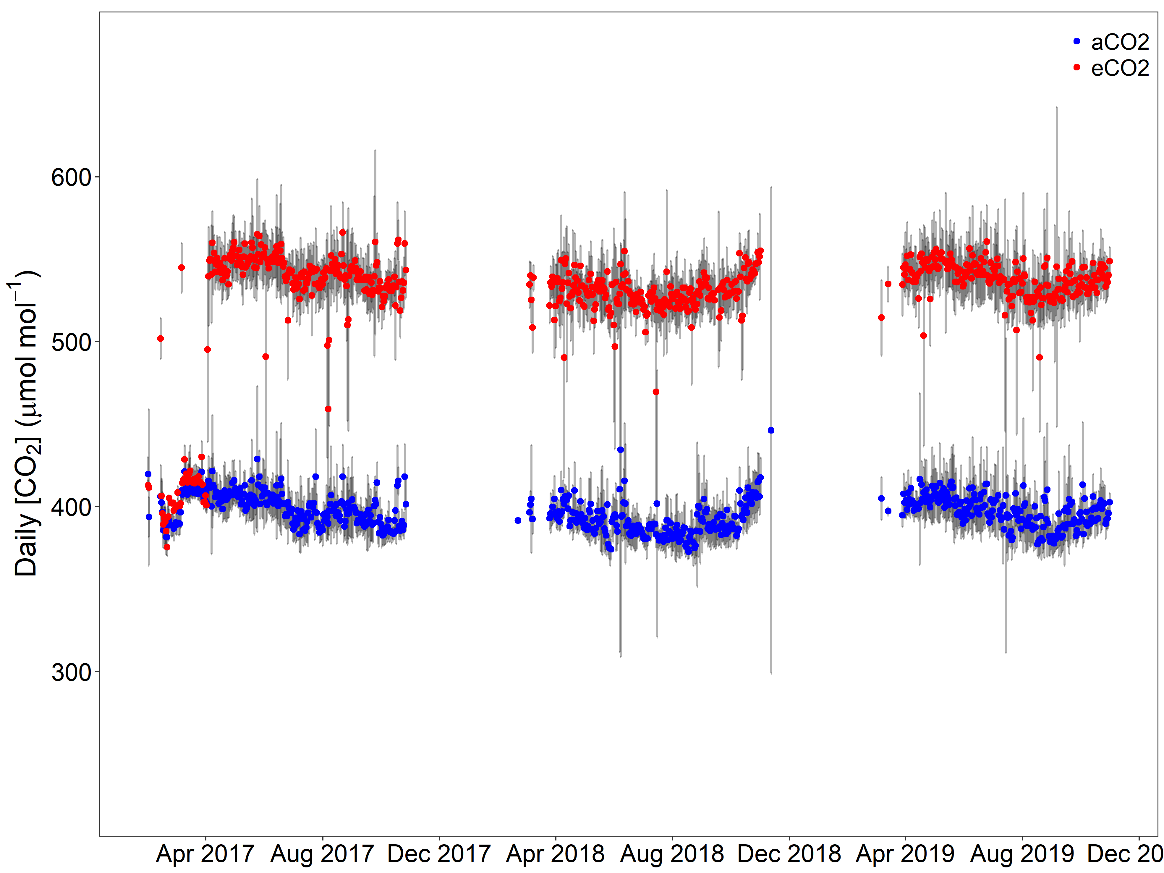

Supplementary Figure 1. Seasonal course of daily average ambient CO_2_ (blue) and daily average elevated CO_2_ (red) at the BIFoR FACE facility. All data are calculated using one minute CO_2_ averages from January 2017 to December 2020. Grey bars shows the standard deviation between the three replicate plots (n=3). Data are shown for all times when the FACE system was scheduled to operate from April to October and therefore includes all engineering failures and automatic shutdowns due to inclement weather.

**Supplementary Figure 2.**

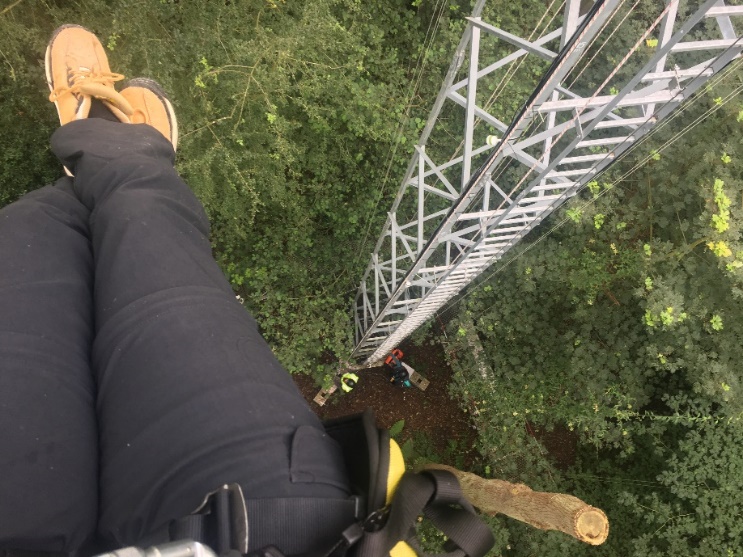

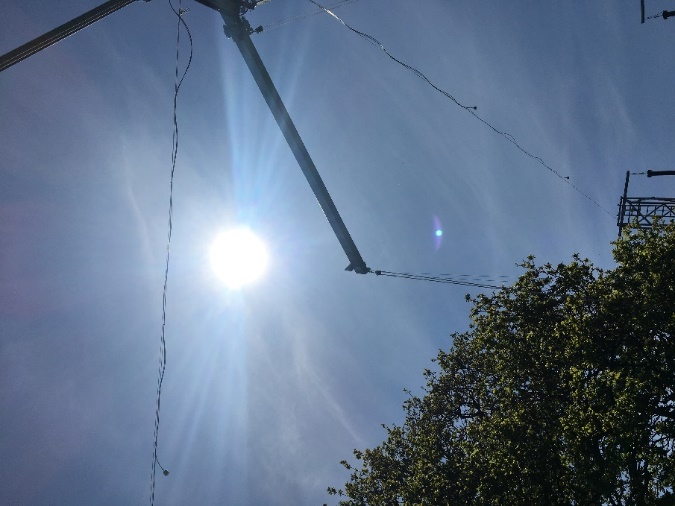

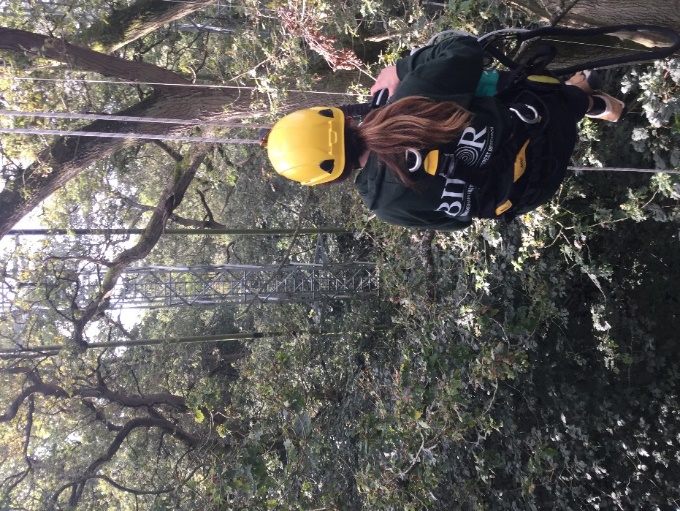

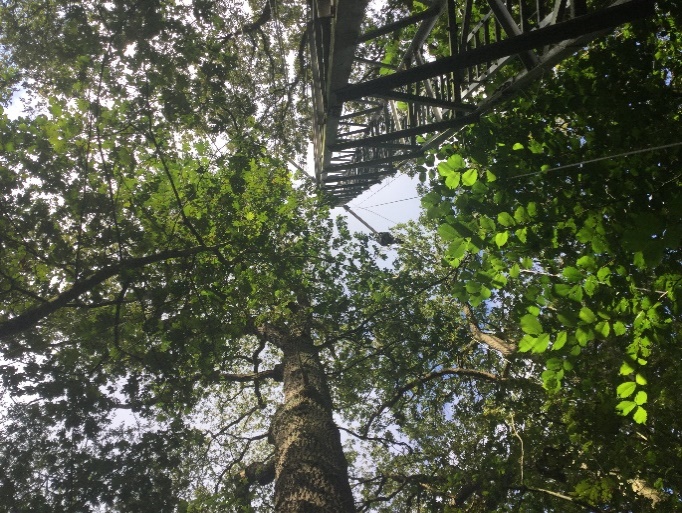

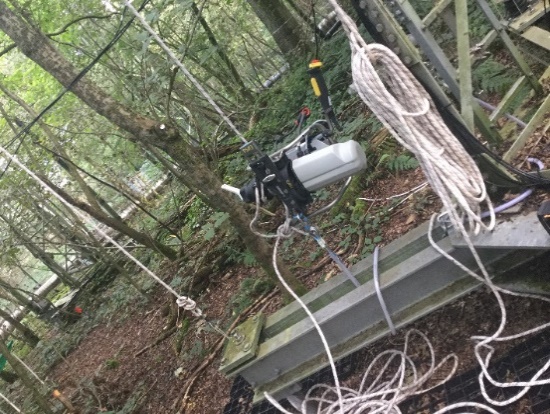

Supplementary Figure 2. Photographs detailing the canopy access system (CAS) at the BIFoR FACE site.

**Supplementary Figure 3.**

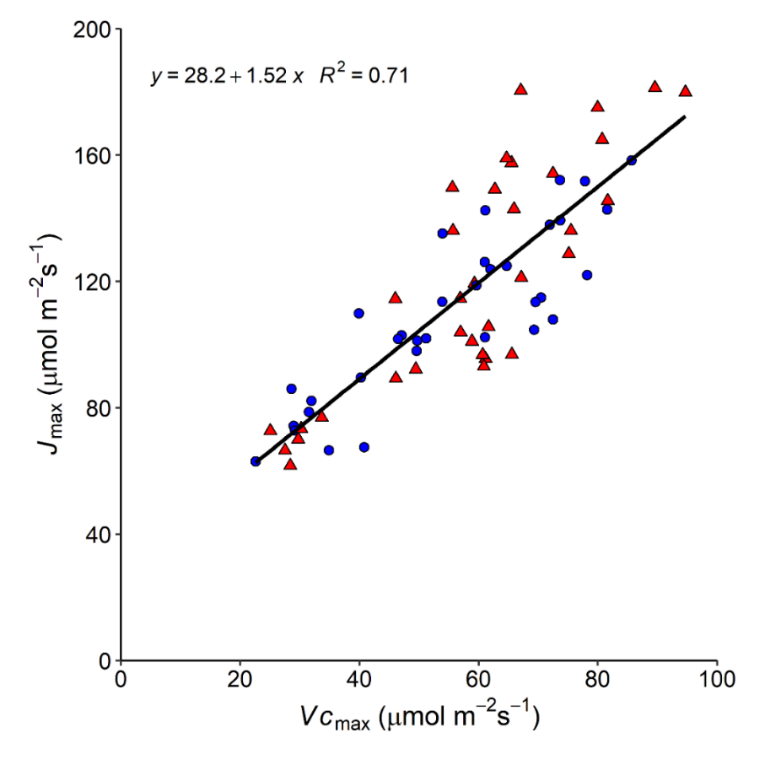

Supplementary Figure 3. Linear relationship between maximum rates of electron transport (*J*_max_) and maximum rate of Rubisco carboxylation (*V*_cmax_) in upper canopy *Q. robur* at BIFoR FACE. *V*_cmax_ and *J*_max_ were derived as best-fit estimates from individual *A—Ci* curves. Data points represent trees that were exposed to either ambient (blue circles) or elevated (red triangles) CO_2_ concentrations in 2018 and 2019. A single regression line is presented because the regressions generated for each treatment were not statistically different. The regression for the combined data was statistically significant (*P <* 0.0001). Individual regressions for ambient, $J_{max}^{a}\left( {Vc}_{max}^{a} \right)$, and elevated, $J_{max}^{e}\left( {Vc}_{max}^{e} \right)$, concentrations are: , $J_{max}^{a}=38.5+1.29 {Vc}_{max}^{a}$ , R^2^ = 0.75 and $J_{max}^{e}=19.1+1.72 {Vc}_{max}^{e}$, R^2^= 0.71.

| **Supplementary Table S1. ANOVA to compare light categories used in Figure 4.** | | | | | |
| --- | --- | --- | --- | --- | --- |
|  | | **Light category** | | | |
|  | | >1000 | >500- <1000 | >250- <500 | <250 |
| Source of variation | Df | *P*-value | *P*-value | *P*-value | *P-value* |
| **CO_2_** | 1 | 0.59 | 0.34 | 0.18 | 0.33 |
| **Year** | 1 | 0.55 | 0.24 | 0.73 | **0.011*** |
| **CO_2_ * Year** | 1 | 0.88 | 0.66 | 0.64 | 0.62 |

| **Supplementary Table S2. ANOVA to compare sampling years in Figure 5.** | | | | | |
| --- | --- | --- | --- | --- | --- |
|  | | ***V*_cmax_** | ***J*_max_** | **N*_m_*** | **N*_a_*** |
| Source of variation | Df | *P*-value | *P*-value | *P*-value | *P*-value |
| **Year** | 1 | 0.099 | 0.23 | 0.21 | 0.09 |

| **Supplementary Table S3. ANOVA to compare sampling years in Figure 6.** | | | |
| --- | --- | --- | --- |
|  | | **A_net_** | **Response ratio** |
| Source of variation | Df | *P*-value | *P*-value |
| **Year** | 1 | 0.103 | 0.41 |

**Supplemental Appendix 1.**

**Calculating a theoretical enhancement of photosynthesis.**

Before starting the BIFoR experiment, we asked what the *a priori* theoretical enhancement of photosynthesis would be for +150ppm over current ambient CO_2_ of 400ppm. There are a variety of approaches to define this that can be taken. For one such approach, Nowak et al. (2004) reasoned that all photosynthetic CO_2_ responses can be collapsed to a common shape, normalising the maximum CO_2_- and light-saturated rate of A_net_ (called A_max_). From the biochemical model of Farquhar et al. (1980), we can compute how far below the A_max_ one could expect ambient and elevated A_net_ measurements could be. Specifically, the elevated A_net_ is expressed as a fraction of the normalised A_max_. The baseline ambient A_net_ can be equally computed. Then it is possible to take the ratio of these percentages below the A_max_ to get a simple, theoretical relative enhancement value.

Following this approach and using a pre-experiment parameterisation of the biochemical model, we used the "Photosyn" package in R to simulate an expected photosynthetic enhancement for *Q. robur* trees at BIFoR. The intent of the simulations was to estimate an approximate A_net_. Photosyn employs a coupled photosynthesis - stomatal conductance model (Duursma, 2015), based on the Farquhar et al. (1980) model of photosynthesis, and an optimisation model of stomatal conductance (see Duursma 2015). The model considers the temperature sensitivity of photosynthetic parameters, and a dark respiration rate calculated from leaf temperature and scaled to day respiration. The parameters we used for the model for *Q. robur* to simulate ambient A_net_ and the other needed rates of photosynthesis were *V*_cmax_ = 48 µmol m^‑2^ s^‑1^ with *J*_max_ slaved to *V*_cmax_ according to a factor of 1.85 appropriate for 25 °C, and saturating light (*Q* = 1800 µmol m^‑2^ s^‑1^). The g1 for the stomatal optimisation slope was 1.5 for the simulations, which yielded a Ci/Ca ratio during gas exchange of 0.7. From this parameter set, the photosynthetic enhancement for a +150 ppm CO_2_ enrichment was 1.35. We also iterated this calculation for alternate *V*_cmax_ (40 to 80 µmol m^‑2^ s^‑1^) and g1 values (1.0 to 2.5), producing photosynthetic enhancement values between 1.35 and 1.39. As a result, in the main portion of this manuscript we used a 1.37 photosynthetic enhancement for the +150 ppm CO_2_ enrichment as a middle value for our theoretical expectation emerging from the model simulations.
